# Cat brains age like humans: Translating Time shows pet cats live to be natural models for human aging

**DOI:** 10.1101/2025.07.31.667772

**Authors:** Capucine Januel, Elijah Morrow, Ryan Gibson, Amanda Gross, Alexandra A. de Sousa, Brier A. Rigby Dames, Christine J. Charvet

**Affiliations:** Department of Anatomy, Physiology & Pharmacology, College of Veterinary Medicine, Auburn University, Auburn, Alabama, USA; Ecole Nationale Vétérinaire de Toulouse, France; Scott-Ritchey-Research Center, College of Veterinary Medicine, Auburn University, Auburn, Alabama, USA; Centre for Accountable, Responsible, and Transparent AI (ART-AI), Department of Computer Science, University of Bath, Bath, Bath, UK

## Abstract

Translating biological time across species is a powerful tool to identify new models of human aging and disease. Currently, it is not clear whether any animal reaches an age comparable to a human in their 80s. Most species seem to age differently compared with humans. Some preliminary observations suggest that cats may share common patterns of aging with humans. Cats could serve as a promising model for human aging. Here, we find corresponding ages between cats, humans and other species to test whether cats can live to the equivalent of a human in their 80s. We analyzed 3,754 observations across species from sudden and gradual changes in anatomy, physiology, and behavior. Some of these data are from clinical records, whereas others are from brain scans using high-resolution MRI (7T and 3T). We studied pet cats, research colony cats, and wildcats living in zoos to encapsulate species variation in the speed of development and aging. We found that cat and human brains atrophy with age, and that their age-related patterns in brain aging are sufficiently similar that we could use them to generate cross-species age alignments. We also found that human postnatal development is stretched compared with cats and mice. Interestingly, some pet cats that visit clinics are much older than those in colonies. Therefore, cats, and especially pet cats, are natural model systems of human aging. Our findings call for increased integration across veterinary and human medicine to understand aging.

## Introduction

Human aging is characterized by cognitive decline, brain atrophy, and increased incidence of geriatric diseases such as Alzheimer’s disease (AD), which is characterized by memory loss and cognitive impairment (Nelson et al., 2009). Age-related changes, including brain atrophy and AD were considered unique to humans because these traits were hard to identify in other species (Finch and Austad, 2015). Mice are the most studied species in aging research, even though they do not spontaneously share many hallmarks and diseases of aging (Yokoyama et al., 2022; Rigby Dames et al., 2023; de Sousa et al., 2023). Broad surveys show that some animals develop brain amyloid pathology and tangles (e.g., cats, chimpanzees; Youssef et al., 2016), suggesting some species recapitulate human aging patterns, and may serve as models for human aging. We test whether cats live sufficiently long to recapitulate age-related changes.

It is an open question as to whether an animal lives sufficiently long to map onto aged humans (e.g., an octogenarian). According to AnAge, the maximum human lifespan is 122.5 years, reaching a far longer lifespan than closely related species (e.g., 68 years in chimpanzees; de Magalhães and Costa, 2009; de Magalhães et al., 2024). Domestic cats also have an extended maximum lifespan (*Felis catus:* 30 years) compared with related species (e.g., wildcat; *Felis silvestris:* 19 years; Tacutu et al., 2013; Teng et al., 2024), and there are an estimated 600 million cats worldwide (Driscoll et al 2009; Griffin and Baker, 2007). In contrast, there are a few thousand captive chimpanzees in the United States (Knight, 2008). Cats may be well-suited to study human aging by virtue of their large numbers and extended lifespan.

Many pet cats develop chronic diseases similar to humans (e.g., cognitive dysfunction syndrome, obesity; Freeman et al., 2006; Sordo et al., 2020). They can also develop amyloid deposits and hyperphosphorylated tau aggregates (Rofina et al., 2006; Landsberg et al., 2012; Chambers et al., 2015, Seibert et al., 2017; Sordo and Gunn-Moore, 2021; Sordo et al., 2021). Cats may live long enough to recapitulate human age-related changes. We characterize feline aging and find corresponding ages across the lifespan of humans and cats.

During aging, the brain atrophies, and many brain structures change. The cerebral cortex thins, cortical sulci widen, ventricles enlarge and the interthalamic adhesion (i.e., a midline structure connecting thalami) thins. Human brain atrophy is a normal part of aging and becomes salient from 50s-60s onwards (Long et al., 2012), and it is exacerbated in dementia-related diseases (Fjell et al., 2017; Coupé et al., 2019; Blinkouskaya and Weickenmeier, 2021). Thus far, age-related changes in brain structure have been found in a handful of animals, including chimpanzees and lemurs (Kraska et al., 2011; Picq et al., 2012), though the extent of brain atrophy appears rather modest (Chen et al., 2013). Indeed, few chimpanzees show brain atrophy, or other age-related changes(e.g.., menopause; Rosen et al., 2008; Chen et al., 2013; Edler et al., 2017). Many aging-related processes occur past our 50s, but chimpanzees do not live long enough to show the full spectrum of these age-related changes because few of them live past their 40s, which equates to humans in their 50s (Charvet, 2021). It is an open question as to whether any animal lives long enough to show human aging patterns.

We used different metrics to align ages across species (e.g., bloodwork; Charvet et al., 2025; Cottam et al., 2025). We used MRIs to test whether humans and cats share common patterns in brain aging. We also studied animals in different environments, and we compared the pace of development between domestic and wildcats. We found that pet cats are studied at older ages than those housed in colonies, cat brains atrophy in a manner similar to humans, and that cats can live to the equivalent of a human octogenarian.

## Materials and Methods

We integrated observations with previously published data to equate ages across species (Figure 1-2; Clancy et al., 2001; Workman et al., 2013; Charvet and Finlay, 2018; Charvet et al., 2022; Cottam et al., 2025). The data set includes 3,754 observations (e.g., age of eye-opening in a cat) from different sources, including age-related changes in brain structure from MRI scans, blood chemistry profiles, bone ossification, behavior, and epidemiological data from cats, humans, chimpanzees, rats, and mice. Whereas an “observation” focuses on a single datum point, “time point” refers to several observations recorded in at least two species (e.g., age of eye-opening). Some time points are separated by sex (e.g., reproduction, body growth), while other time points may be drawn from both males and females. Our approach permits considering multiple observations per time point in each species. We recorded distributions from observations (e.g., minimum age of eye opening) and we included different statistics for a given time point in these comparative analyses. We distinguished captive versus wild animals when the information was available. We warn against over-interpreting subtle differences across groups because observations are from multiple sources and researchers may use slightly different definitions across studies. We used Web Plot digitizer v. 4 (Rohatgi, 2025) to extract some data from published plots. All statistical analyses were performed with the programming language R.

**Figure 1.**
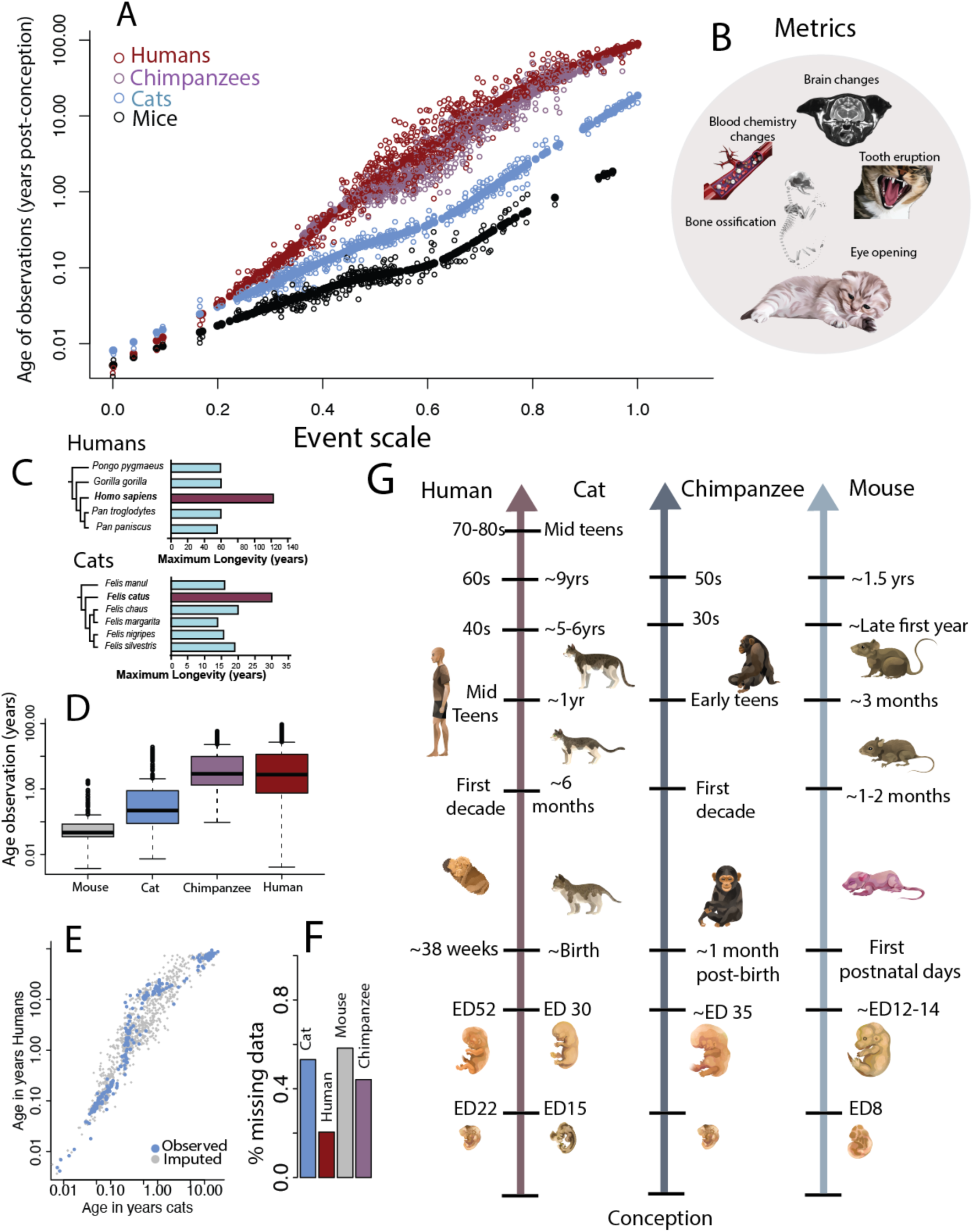
We used a linear model with a spline to translate ages across species. (A) Time points expressed in years post-conception are plotted against the event scale. The smooth spline allows for variation in the pace of development and aging. (B) We used a range of metrics to align ages. Those include age-related changes in brain structure, blood work, the timetable of bone ossification, and behavioral milestones such as eye opening. (C) Cats, like humans, have a prolonged duration of maximum lifespan compared with closely related species. (D) Age ranges of observations span much of the known lifespan of humans and other species. (E) We imputed observations when not available in some species with an example shown for cats and humans. (F) Percentage of missing data are shown for each species. (G) We provide some examples of age alignments generated from the model. Abbreviations: PM: postnatal month; P: postnatal day.

**Figure 2.**
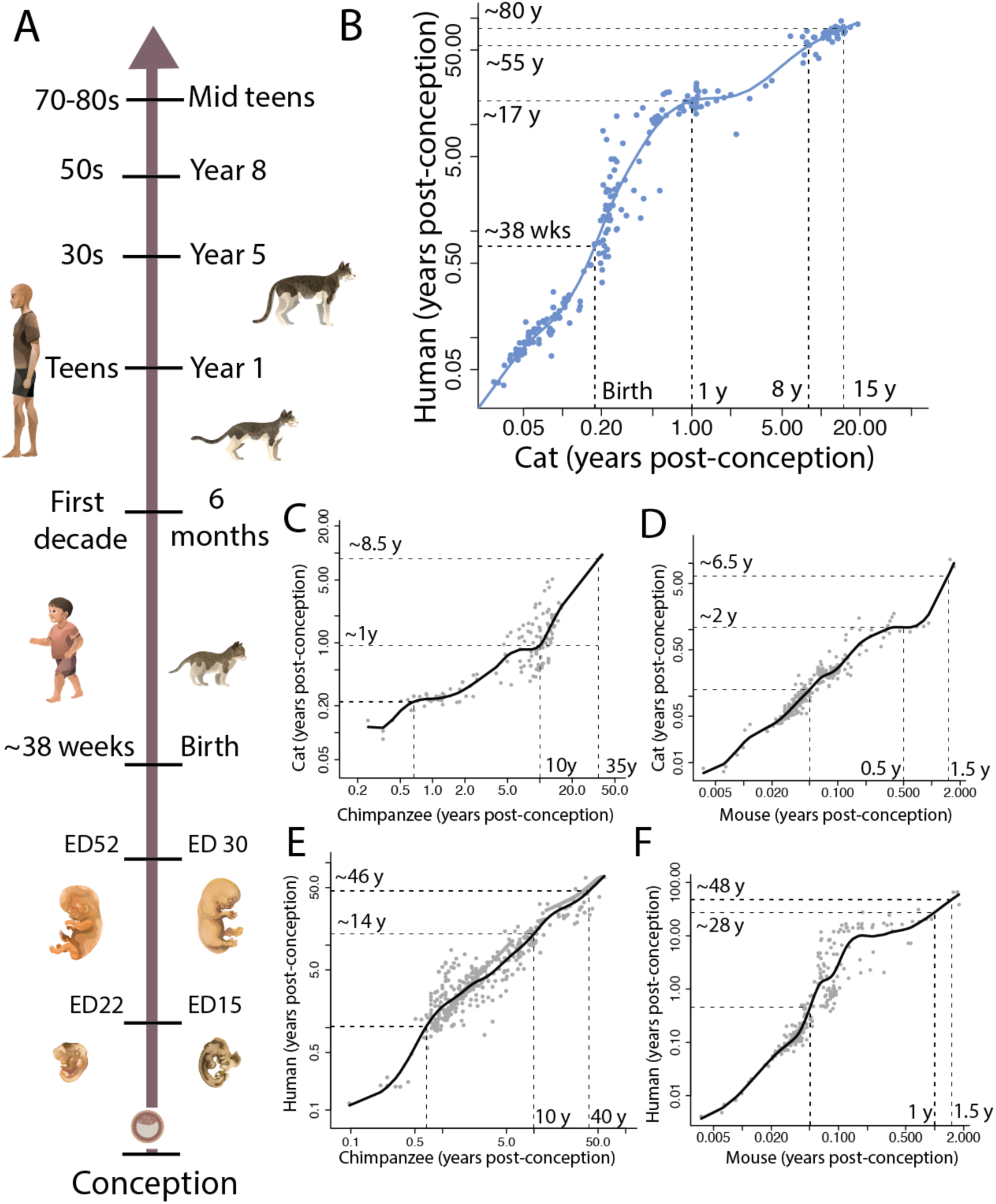
(A) Example age matches between humans and cats. A newborn cat equates to a human at 40 weeks post-conception, and a 6-month-old cat equates to a human within their first decade, and a cat in their teens maps onto a human in their 80s. (B-F) Scatterplots of two species comparisons demonstrate that there is a complex relationship in the pace of development and aging across species. Dashed lines show examples of corresponding ages between two species.

### Brain necropsy reports

Brain necropsies are not typically performed in cats lacking neurological signs, but we found 4 cases reporting brain atrophy from cat necropsy reports from the Auburn College of Veterinary Medicine. These necropsies mentioned cerebrocortical atrophy, hippocampal atrophy, and widening of cortical sulci, suggesting that aged cats show brain atrophy. These post-mortem mentions guided our analyses from *in vivo* MRI scans (e.g., whole brain volumes, gyrification, Figures 3-5).

**Figure 3.**
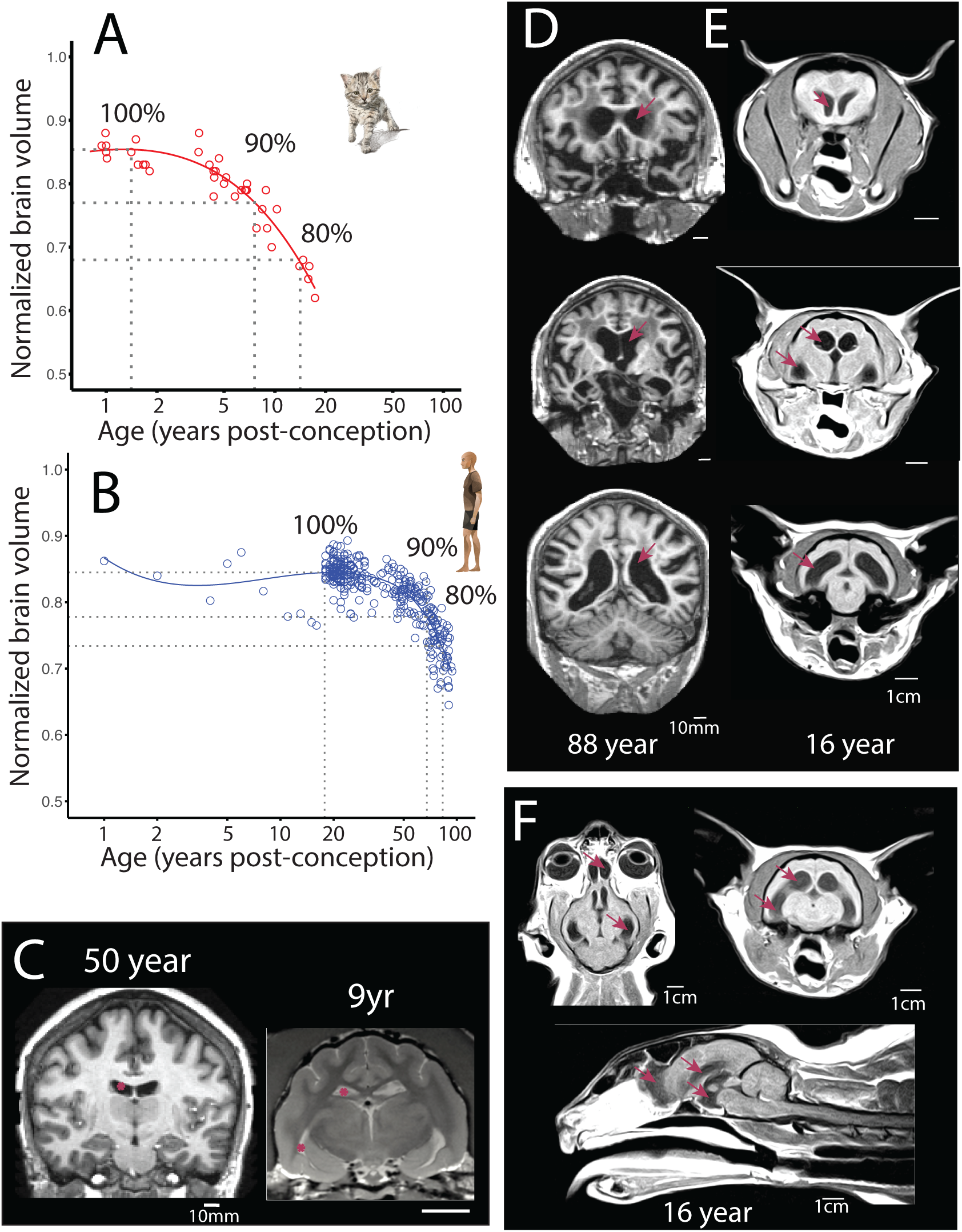
The brain atrophies with age in (A) cats and (B) humans. We fit a smooth spline to multiple metrics (e.g., normalized brain volume) versus age to extract corresponding observations across cats and humans. We, for example, extracted the age of peak brain volume (100%) before the brain begins to decline and the age at which the volume reaches a particular percentage of adult brain volume (e.g., 90, 80% of maximum brain volume). We use these time points to generate cross-species age translations. (C). A 50-year-old human (C) and a 9-year-old cat (C) begins to show modest enlargement of ventricles (red asterisks). At later ages, the ventricles are much enlarged (arrows) as exemplified in an 88-year-old human (D) and a 16-year-old cat (E). (F) Horizontal, coronal and sagittal slices show the extent of the brain atrophy in the same 16-year-old cat. In this cat, we found atrophy of the olfactory bulb and telencephalon in addition to an expansion in lateral and third ventricles (arrows).

### Cat brain MRI

We used multiple MRIs machines (3T and 7T MRI scanners) to study brain structure in cats housed at Auburn University and from patients in the veterinary clinic. These MRIs were collected for purposes other than the present study, which obviates the need for IACUC approvals for the purpose of this study. Cats from the Auburn veterinary clinic varied from 4.33 months to 17.3 years of age post-birth (n=18; 12 males, 6 females). A board-certified veterinary neurologist evaluated health records and MRIs to exclude cats with brain lesions. Colony cats receiving MR scans were significantly younger than those visiting the clinic. The ages of cats from the colony varied from 0.07 to 10.29 years after birth (n=34; 10 males, 24 females; mean age: 2.53 years after birth). Whereas the ages of healthy cats from the colony are well-documented, the age of pet cats visiting the clinics are supplied by the owner and these cats were scanned for health issues (e.g., seizures), which may impact brain measurements. We therefore compared brain structure metrics between colony and pet cats to evaluate whether health status and age uncertainty impact measurements.

### Cognition

We did not find evidence that cats in our study have cognitive dysfunction syndrome. We considered behavioral changes, such as spatial or temporal disorientation, vocalizations, changes in interactions with others and sleep-wake cycles, activity levels, house-soiling, anxiety, learning and memory deficits in these cats (Sordo et al., 2020) even though we do not have a recognized validated diagnostic tool to evaluate dementia in cats. Researchers overseeing the cat colony did not observe clear signs of cognitive dysfunction. A board-certified veterinary neurologist evaluated the health records of pet cats receiving MRIs in the clinic and confirmed a lack of evidence these cats have cognitive dysfunction syndrome.

### 7T MRI scanner

We used the 7T MRI Siemens MAGNETOM clinical scanner. Cats were anesthetized and scanned as part of other long-term efforts to study lysosomal storage disease (Gray-Edwards et al., 2014). MRI scans were collected with a 28ch Rx x 1ch Tx knee coil (QED). Multiple MRI pulse sequences were collected. These include MPRAGE (Magnetization Prepared Rapid Acquisition Gradient Echo), SPACE (Sampling Perfection with Application optimized Contrasts using different flip angle Evolution), T2 TSE (Turbo Spin Echo), HASTE (Half-Fourier Acquisition Single-shot Turbo Spin Echo) and CEST (Chemical Exchange Saturation Transfer). We used either T1 or T2 *in vivo* MR images with a variable TR and TE (e.g., TR=50ms, TE=4ms; TR=5,450ms, TE=12ms). Voxel resolution varied across scans. Some scans were anisotropic (e.g., 0.336mm x 0.336mm x 1.2mm) whereas other scans were isotropic (e.g., 0.5mm x 0.5mm x 0.5mm).

### 3T Clinical MR scanner

Veterinary radiologists and neurologists collected MRI scans of cat brains in the clinic. These cats were anesthetized and scanned with a 3T MRI Siemens Skyra MRI Scanner. The standard procedure is to administer pre-anesthetic medication followed by induction and gas anesthesia with isoflurane before cats are scanned. Scan resolution (anisotropic resolution: e.g., 0.469mm x 0.469mm x 3.3mm; 0.781mm x 0.781mm x 5mm) and imaging parameters (e.g., TR=600, TE=11.5; TR=2,000, TE=10) varied across scans. During these sessions, T1 and/or T2 scans were collected. We used the modality with the highest image resolution for our morphometric analyses. A board-certified neurologist selected MRI scans from the clinic. Cats with brain lesions or brain masses were excluded from the study.

### Human brain scans

We used structural MRI scans of human brains (age range: 1-99 years old) from multiple databases, including OASIS1 (MPRAGE sequence; anisotropic resolution; 1mm x 1mm x 1.25mm), Brain Chart, and NITRC (Bethlehem et al., 2022; Richards, 2019). Details of T1- and T2/FLAIR-weighted structural MR scans are described previously (Marcus, 2007; Bethlehem et al., 2022; Richards, 2019; Fotenos et al., 2005). We used brain measurements from individuals with no evidence of cognitive loss. We used the Clinical Dementia Rating (CDR) score (Berg, 1988), which is a measure of dementia across six cognitive domains (memory, orientation, judgment, problem-solving, community affairs, home and hobbies, personal care). We selected individuals with a CDR score of 0, which indicates a lack of evidence for dementia.

### Volumetric analyses

We collected and we used past measurements from human and cat brain scans (Figures 3, 4). Figure 4 shows example measurements of some brain metrics. We contoured the brain, the cranial box, and the lateral ventricle from brain slices to reconstruct the volume. The brain volume measurements did not include the lateral ventricles. The cranial box volume was calculated by subtracting the brain from the cranial volumes. Normalized whole brain volumes (nWBV) was computed by dividing brain volume by the cranial box. We used published data for the whole brain and cranial box volumes from the OASIS dataset (n=436: Marcus, 2007), which segmented the brain into cerebrospinal fluid, gray matter, and white matter via a hidden Markov random field model. We used human lateral ventricle volumes from the Brain Chart (Bethlehem et al., 2022). We used Image J and OsiriX to quantify changes in brain structure in humans and in cats.

**Figure 4.**
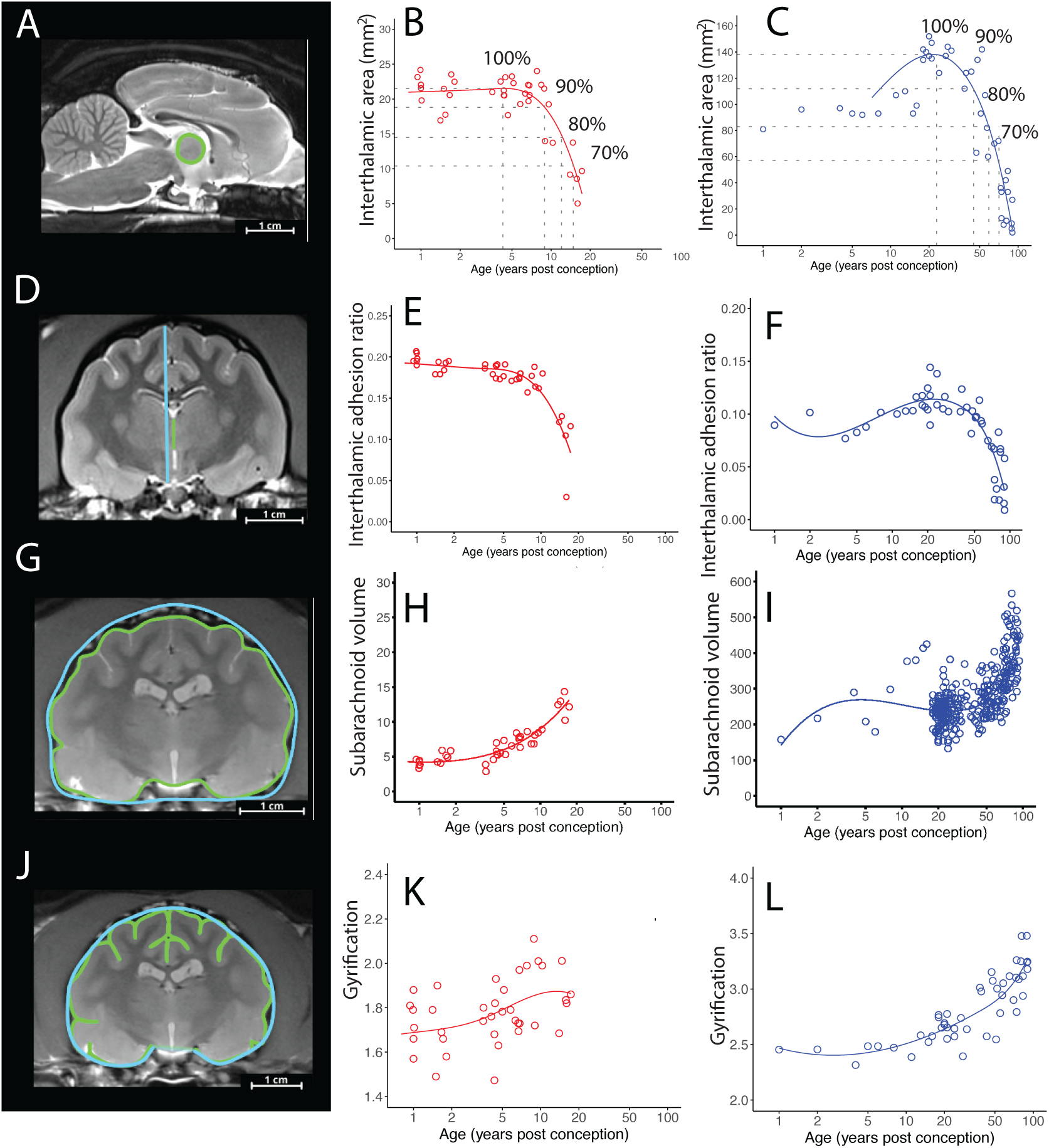
Sagittal and coronal slices from brain scans from the 7T MRI illustrate some of the brain measurements collected from colony cats. (A) We used sagittal slices to measure the interthalamic area (color-contoured). The interthalamic adhesion area peaks and declines with age in (B) cats and (C) humans. We fit smooth splines to capture ages of peak interthalamic adhesion area before it declines to specific percentages (e.g., 90%, 80%, 70%) of its maximum value (B-C). We used coronal slices to measure the interthalamic adhesion relative to the brain height (D-F) measured along the brain’s midline. The interthalamic adhesion ratio declines with age in (E) cats and (F) humans. (G) We also measured the cranial space by quantifying the volume of the brain and cranial box. To do this, we measured the brain and cranial box (color-contoured; G) from coronal slices. The intracranial space represented as subarachnoid volume increases drastically in (H) cats and (I) humans. (J) We quantified gyrification as the ratio between the perimeter contouring gyri and sulci relative to the perimeter contouring the brain. Gyrification increases with age in (K) cats and (L) humans. The cats shown above are 4.45 years old (A), and 8.85 years post-birth (D, G, J).

**Figure 5.**
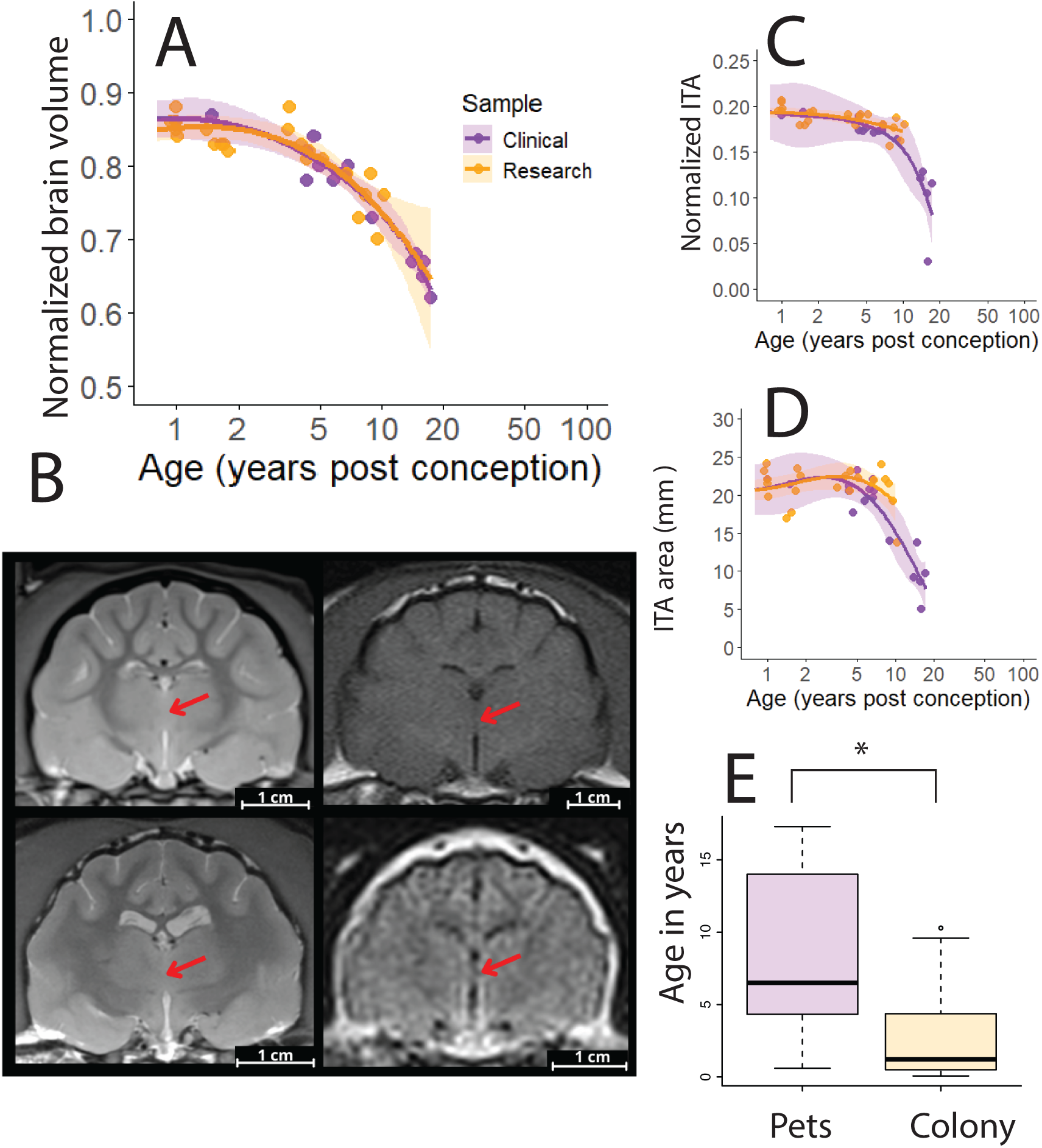
(A) Normalized brain volume declines with age in both colony and pet cats. We fit smooth splines and their confidence intervals to Auburn colony and clinical cats separately and we found extensive overlap between these two feline populations even though we used different MR scanners (B) to collect these brain scans. (B) Coronal slices show example images collected with the 7T MRI scanners (left panels) and the 3T MRI scanners (right panels). The ages of study varied (top left; 0.62 y; bottom left: 8.85 y; top right: 4.66 y; bottom right: 17 y). Colony cats were scanned in a 7T MRI which can generate scans of higher resolution than pets visiting the clinic. Bran metrics, including the normalized interthalamic adhesion thickness (C) and the interthalamic adhesion area (D) are also very similar across the two groups of cats. (E) Pet cats visiting the clinic are significantly older than those being studied in the laboratory. We used the highest resolution MR imaging modality. This meant we used T1 in some cases or T2-weighted images in other cases. Abbreviation: y-years after birth.

### Brain region metrics

We collected several metrics from the brains of cats and humans to track age-related changes in brain structure. We measured the interthalamic adhesion, which is a bridge of tissue that connects both thalami as well as gyrification. The interthalamic adhesion is a marker for brain atrophy in companion animals (Noh et al., 2017), which varies with age in humans and companion animals (Figure 4; Noh et al., 2017; Blinkouskaya and Weickenmeier, 2021; Hasegaw, et al., 2005). We used a coronal (i.e., transverse) slice to measure the interthalamic adhesion thickness (where thickest) and the brain height. The brain height is defined as a vertical distance from the highest dorsal point to the most ventral point. We considered age-related changes in interthalamic adhesion thickness and normalized it to the brain’s height. We also measured the interthalamic adhesion area from a sagittal slice, which was along the brain’s midline where the thalami are smallest. We used a coronal slice to define gyrification as the ratio between the perimeter contouring gyri and sulci relative to the perimeter of the brain’s contour. We measured these perimeters from the same coronal slice used to measure the interthalamic adhesion thickness. Table S1 lists details about brain measurements.

### Brain metrics to align age

For each species, we fit a smooth spline to model the relationship between age and brain structure metrics, such as interthalamic adhesion thickness and normalized brain volume. We identified the age at which brain metrics (e.g.., interthalamic adhesion, brain volume) are at their maximum or its minimum before they decline or increase with age. If the metric decreased with age, we used the smooth spline to calculate the age at which the values fell to specific percentages (e.g., 90%, 80%, 70%). We fit smooth splines separately by sex (male versus female cats) to analyze sex-related differences and by population (colony versus clinic cats) to evaluate whether MRI or health status might impact brain structure metrics. These analyses are based on 53 cats (22 males, 31 females) and 446 human participants (n=171 males, n=275 females).

### Health records to equate ages across species

Many metrics are collected for health purposes, and these metrics vary with age. For example, alkaline phosphatase (ALP) declines postnatally and subsequently becomes relatively invariant. These age-related patterns are conserved across different mammalian species (e.g., cats, chimpanzees, humans). Similarly, body weights increase postnatally, plateau, and subsequently decline later in life. If age-related changes are similar across studied species, we fit non-linear regressions to patterns of change common across species to extrapolate corresponding ages across populations with the library R package easynls (e.g. model=3). For example, we first fit a smooth spline through ALP versus age and we extrapolated corresponding values from the smooth spline. We then used the R package easynls to test whether a piecewise linear model (y ∼ a + b * (x - c) * (x <= c)) accounts for a significant percentage of the variance. We used this model to extrapolate the age the plateau is reached. We also quantified when the values reached a percentage of the plateau (e.g., 120% of the plateau). We use these as observations in the model used to equate ages across species. We used health records from multiple sources, including the Auburn Veterinary Clinic, the colony, Project CatAge, and the primate aging database (PAD). We expect data from PAD and the colony to come from healthy animals. The health records from pet cats visiting the clinic and from PAD were not re-evaluated by a clinical pathologist, as was the case for colony cats.

### Colony cats’ blood chemistry profiles

Blood chemistry profiles from domestic shorthairs in the colony were collected from 2016 through 2021. A board-certified veterinary clinical pathologist re-evaluated health metrics collected from SRRC colony cats to confirm that the cats were healthy (n=99 samples; n=52 individuals; min-max: 0.12-9.9 years after birth; mean: 2.6 years after birth). These data were collected for purposes other than this study.

### Auburn CVM blood chemistry profiles

We used blood chemistry profiles from apparently healthy cats collected from Auburn clinic from 2015 to 2023 (e.g., n=224 ALP observations; 197 cats; 85 females and 84 males; range min-max: 0.39-17.82 years after birth; mean: 8 years after birth). Most of these chemistry profiles are from domestic shorthairs, a few mixed breeds, some domestic longhair cats, and a handful of specialty breeds (e.g., Siamese cats, n=5; Ragdoll cats, n=3). These health metrics were not collected for the purposes of this study, which obviates the need for IACUC approval. All information about the cat’s owner was removed before we gained data access.

### Project CatAge blood work

We asked cat owners to provide information about their cat’s health records and life history (e.g., age, indoor or outdoor) on social media. We included cats if their owners are confident of their cat’s age or if they obtained them when they were kittens (i.e., less than 8 weeks of age). Project CatAge was IRB exempt by Auburn University. We collected 91 blood tests (i.e., total protein, ALP, creatinine, phosphorus) from 47 spayed or neutered cats. Some tests do not include all metrics of interest. We excluded 13 blood tests from 10 cats, and we excluded ALP values from two cats for health reasons. We used 78 blood tests from 45 cats (22 males; 23 females; min age: 0.34 years old; max age: 18.12 years old). Most of the cats are domestic shorthairs and live indoors (Figure 6). The cats are from multiple US states, though most are from Alabama and other southern states. The health of these cats had been re-evaluated under the guidance of a board-certified veterinarian.

**Figure 6.**
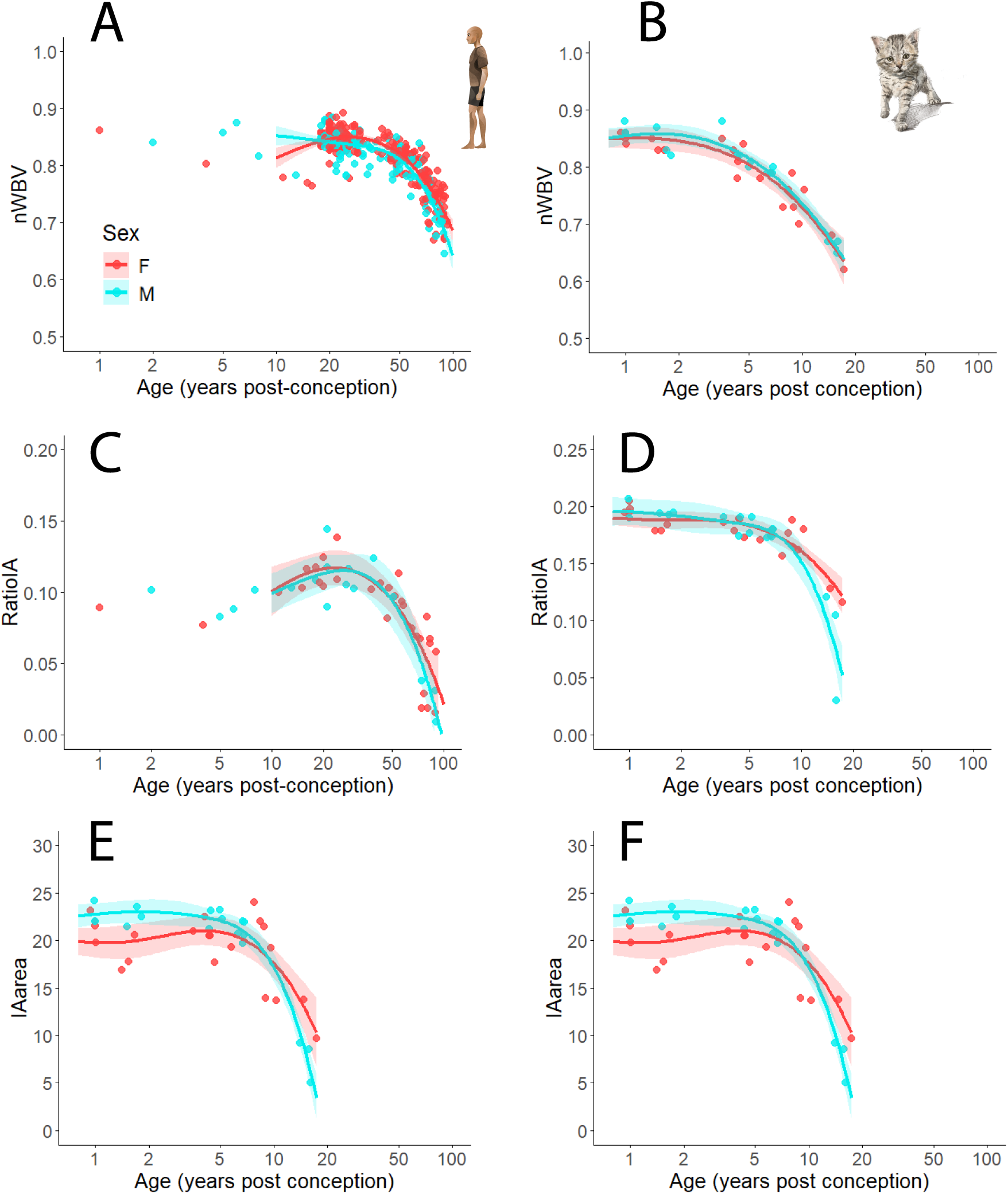
We fit smooth splines and confidence intervals separately for males and females to assess whether there are sex differences in age-related patterns of brain change. Normalized brain volume in humans (A) and cats (B) decrease with age. There is little variation between males and females. The normalized interthalamic adhesion thickness (C, D) and area (E, F) decline with age. Here, there is extensive overlap in normalized brain volume in males and females. Males appear to show accelerated reductions in interthalamic adhesion area and thickness after 10 years old in cats, and after 50 years old in humans.

We included animals’ blood values found within the normal range supplied by each health report, though we included elevated ALP and phosphorus in cats under 1 year of age because these values are typically high in kittens (Levy et al., 2006; Pineada et al., 2013). We also included ALP below the normal range because it is not considered clinically significant (Oikonomidis and Milne, 2023). We excluded blood work from cats with dehydration (Ata et al., 2018; Ford and Mazzaferro, 2011), hyperthyroidism (Vaske et al., 2016), chronic kidney disease (Vervloet et al., 2017), and bladder inflammation (Vaden et al., 2004), as well as those receiving steroids (Ginel et al., 2002) and non-steroidal anti-inflammatory drugs (Hadjicharalambous et al., 2021), because these conditions and medications may impact measured variables. One cat was deemed dehydrated based on several factors, including elevated urine specific gravity beyond the normal range. We excluded three cats once they showed elevated thyroid hormone levels and were diagnosed with hyperthyroidism. We excluded cats with early signs or blood changes associated with chronic kidney disease, but we retained blood values for cats prior to these diagnoses (Syme et al. 2002). One cat was omitted because it had blood in its urine and elevated urine white blood cell counts, which indicates bladder inflammation. Cats with chronic localized conditions (e.g., megacolon, dental disease, osteoarthritis) were included in our dataset if there was no evidence of systemic inflammation or changes to blood parameters (Bhat et al., 2016). The analyses in the present study are based on 78 blood tests from 45 cats.

### Neuropathologies

We collated information on the minimum age of reported Alzheimer-related neuropathologies (e.g., brain amyloid, tangles) across studies (de Sousa et al., 2023) for humans, chimpanzees, and cats (deSousa et al., 2023). We excluded individuals with neurological conditions associated with early AD pathology (e.g., Down syndrome; see Gunn-Moore et al., 2007; Davidson et al., 2018), but we included late-onset seizures, defined as occurring after 5 years of age in cats and after 60 in humans (Hazenfratz and Taylor, 2018; Cleary et al., 2004), because AD-related pathology may have a link to seizures (Sordo and Gunn-Moore, 2021).

### The translating time model

We selected time points with observations found in at least two species. The dataset contains missing data (Fig. 1). We used the library package Amelia to generate imputed datasets (n=10). Imputed values could not exceed the maximum reported lifespan for the relevant species (Fig. 1B). We selected the dataset generating the highest minimum correlation in observations across individuals. We then denoised the imputed dataset with a principal component analysis. We used the imputed dataset to calculate an event scale (example in Fig. 1E), which is an ordering of time points averaged across species (Fig. 1A). We produced a weighted average inversely proportional to the amount of missing data per species (Fig. 1F), which assumed that the least complete data should factor less in the calculation of the event scale. The event scale is calcuated by subtracting each averaged time point by the minimum time point and dividing that value by the difference between the maximum and minimum time point. Accordingly, the event scale ranges from 0 to 1. We then fit a linear model with a natural spline to accommodate selective accelerations and decelerations in the pace of development and aging.

## Results

We first compare age-related variation in brain structure in humans and cats, and we used observations across multiple cat populations to compare the pace of pet, colony and wild cats. We use these data to generate a linear model that equates ages across the lifespan of humans, cats, and other species (i.e., chimpanzees, mice).

### Cats and humans share age-related changes in brain structure

We studied age-related changes in brain structure in cats and humans. Brain atrophy becomes more pronounced with age in cats and humans (Figure 3A and B). Cats in their teens and humans in their 80s possess relatively enlarged ventricles and a concomitant reduction in brain volume as is evident in an 88-year-old human and a 16-year-old cat (Figure 3D-F). We fit significant smooth splines to brain metrics as a function of age to compare age-related trajectories in humans and cats (see Table 1 for significance results). We found that normalized brain volume is rather invariant at early ages and that it subsequently declines gradually. For example, there is a 10 percent decrease in normalized volume between 5 and 10 years of age in cats (Figure 3A) and after 50 years of age in humans (Figure 3B). Some metrics decrease whereas others increase with age. In humans, as in cats, the interthalamic area and the relative interthalamic adhesion thickness decrease with age whereas the subarachnoid volume and gyrification increase with age (Figure 4). These findings demonstrate that brain atrophy shares a common pattern in humans and cats.

**Table 1.** The summary statistics from smooth splines fit to brain metrics versus age in cats and humans.

| **Metric** | **R<sup>2</sup>** | **Adjusted R<sup>2</sup>** | **p-value** | **Species** |
| --- | --- | --- | --- | --- |
| **Interthalamic**<br>**adhesion ratio** | 0.7701 | 0.7492 | 1.615e-13 | Cat |
| **Interthalamic**<br>**adhesion area** | 0.7111 | 0.6842 | 4.152e-11 | Cat |
| **Normalized brain**<br>**volume** | 0.8391 | 0.8245 | < 2.2e-16 | Cat |
| **Subarachnoid**<br>**volume** | 0.8404 | 0.8259 | < 2.2e-16 | Cat |
| **Lateral Ventricle**<br>**Volume** | 0.4939 | 0.4479 | 3.697e-06 | Cat |
| **Gyrus depth** | 0.5073 | 0.4615 | 2.926e-06 | Cat |
| **Interthalamic**<br>**adhesion ratio** | 0.757 | 0.7339 | 2.116e-12 | Human |
| **Interthalamic**<br>**adhesion area** | 0.8155 | 0.7979 | 6.973e-15 | Human |
| **Normalized brain**<br>**volume** | 0.7488 | 0.7458 | < 2.2e-16 | Human |
| **Subarachnoid**<br>**volume** | 0.5282 | 0.5227 | < 2.2e-16 | Human |
| **Lateral Ventricle**<br>**Volume** | 0.9984 | 0.9983 | < 2.2e-16 | Human |
| **Gyrus depth** | 0.755 | 0.7316 | 2.516e-12 | Human |

### Pet cats show brain atrophy and they are older than colony cats

We evaluated whether sex and environment (i.e, pet versus colony) account for variation in patterns of brain atrophy in cats (Figure 5A-D, S1). Comparisons between pet and colony cats is important given differences in image resolution across MRI scanners (Figure 5B). We fit smooth spline models separately for pet and colony cats to examine age-related changes in normalized brain volume (Figure 5A) and interthalamic adhesion measurements (Figure 5C-D). We found that patterns of brain atrophy in pet and colony cats overlapped substantially, as evidenced by the smooth splines found within each other’s 95% confidence intervals. We found significant differences in the age of colony (x=2.77 years old; n=18) versus pet cats (x=7.81 years old; n=34; Kolmogorov–Smirnov test: D test=0.569, p=<0.01, n=18, n=34), which show that pet cats are scanned at older ages than colony cats (Figure 5E). As a result of their extended age, pet cats show more age-related changes in normalized brain volumes (Figure 5A) and interthalamic adhesion measurements (Figure 5C-D) than cats housed in colonies. We also fit smooth splines and 95% confidence intervals to morphometric traits versus age in males as well as in females (Figure 6) and we found data to overlap extensively between males and females in cats and humans, except for the interthalamic adhesion area and thickness towards later stages of life. Both cat and human males show more rapid changes in interthalamic adhesion area and relative thickness compared with females, as is evident from the observation that males fall outside the 95% confidence intervals derived for female data. Our findings highlight possible sex differences in aging patterns in cats in a manner that resembles patterns found in humans. One major finding is that pet cats may be better suited to study aging because they are studied at older ages than those housed in colonies.

### Health metrics show somatic maturation proceeds more slowly in pet than in colony cats

We used clinical metrics to extract corresponding time points across different cat populations (Figure 7A-C) because some of these metrics vary with age (e.g., body weight, blood chemistry profiles). For example, ALP declines postnatally and becomes relatively invariant in different groups of cats (e.g., Auburn clinic, CatAge, colony; Figure 7C). We fit smooth splines at select ages and we applied non-linear regressions to these data to extract the age of plateau as well as the ages at which specific percentages of that plateau value are attained (Figure 7C). We used these metrics (i.e., ALP, total protein, phosphorus, body weights) to compare the pace of development across cat populations (i.e., Auburn clinic, Project CatAge, a colony from Japan and from Auburn; Figure 7E-F). We fit a general linear model with observations from the colony as the dependent variable and those from pets as the independent variable. We integrated all pairwise time point comparisons between colony and pets. This model accounted for a significant proportion of the variance (F=339.5, R² = 0.76, *p*<0.01). We found the slope of the model (y=1.77x-0.23, adj R^2^=0.76; df=104; p<0.01) to be significantly greater than 1 (slope=1.77; t=7.99; p<0.01), which indicates that pet cats mature more slowly than colony cats. Pairwise comparisons between groups consistently show that pet cats mature more slowly than those housed in colonies.

**Figure 7.**
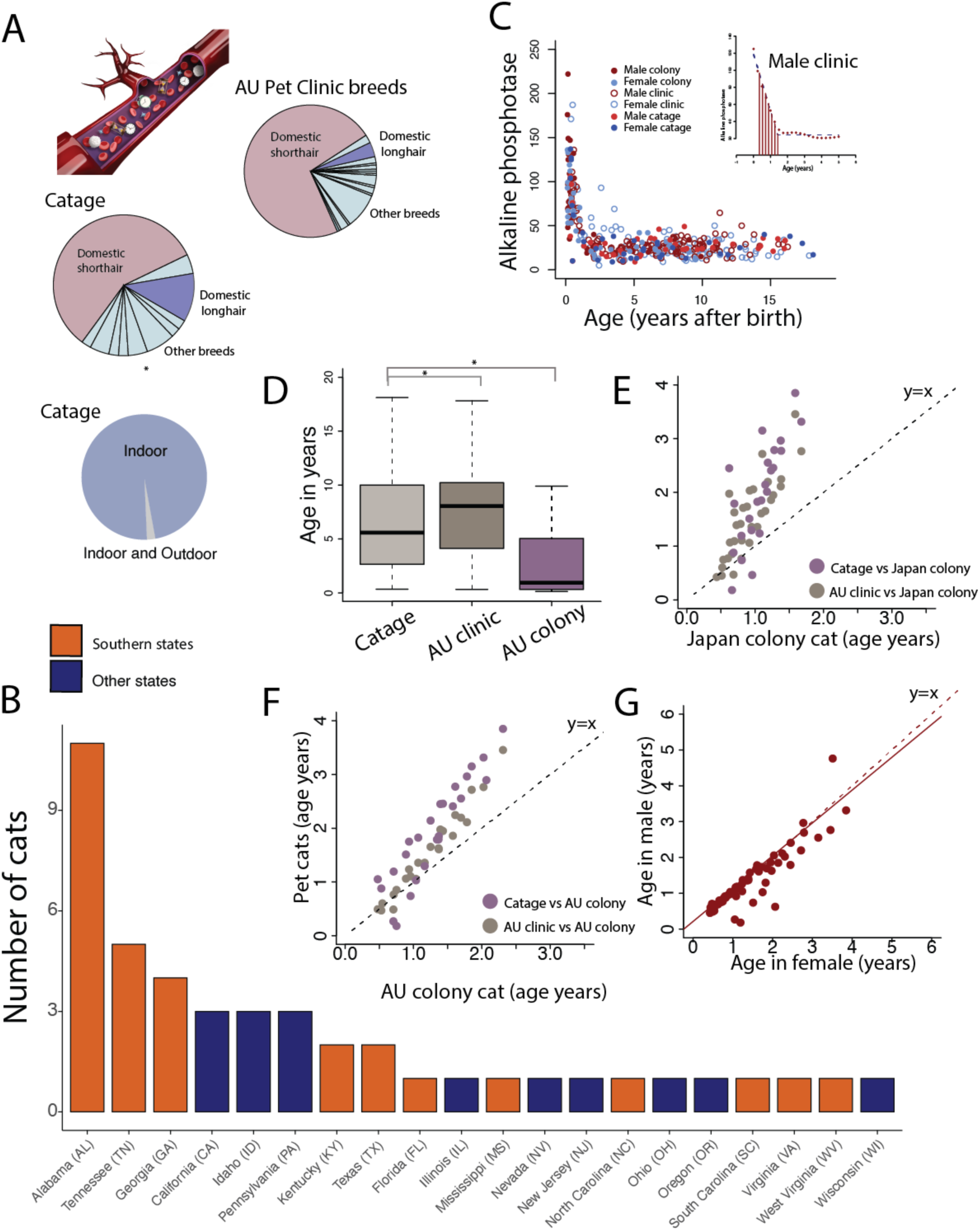
Comparing time points from blood chemistry profiles show that pet cats take longer to mature than those in the colony. (A) Most of the cats used in this study are domestic shorthairs. (B) Cats from Project CatAge are primarily from southern states, but the dataset includes representations across the USA. (C) ALP varies with age and shows common patterns in cats from the colony, the AU clinic, and Project CatAge. We first fit a smooth spline through these data followed by a non-linear model (see inset) to extract the age of plateau as well as epochs (e.g., 90 percent of plateau) within multiple cat groups, including colony cats from Japan and Auburn (Nakai et al., 1992). (D) Cats from the AU clinic (n=169) and Project CatAge (n=45) are significantly older than the colony cats (n=99). Here, we applied a non-parametric Welch one way test followed by a Games-Howell Test to the age of cats for which ALP was measured. An asterisk denotes the pair-wise tests were significant (p<0.01). (E-G) Comparative analysis in time points extracted from blood work values show that pet cats take longer to mature than colony cats. (E) This is evident when comparing all pets to all colony cats though there is no effect for (G) sex. Pairwise comparisons between pets versus the (E) Japan colony and the (F) AU colony consistently show that pet cats take longer to mature than colony cats.

We tested for sex differences in the pace of development amongst pet and colony cats. This is evident given that we fit a general linear model with observations in females (as the dependent variable) versus those in males (as the independent variable) to test for sex differences in the pace of development. The slope of this model (F=239.8, df=59, adj R^2^=0.80; p<0.01, n=61) was not significantly different from 1 (slope=0.92; t value=-1.41; p=0.16; Fig. 7D). These findings indicate that there are no significant differences in the pace of development between males and females.

### Pets with health checks are older than those from the colony

One notable observation from the analysis of feline health records is that studied pet cats are significantly older than colony cats (Figure 7D). Pets with health records have ages that extend to mid to late teens, but this is not the case for colony cats (Figure 7B). We applied a non-parametric Welch test on the age of pet and colony cats from which ALP was measured (F=76.235, p<0.01), followed by a Games-Howell Test for pairwise comparisons to test for significant differences between pet and colony cats (p<0.01). Studied pet cats are significantly older than those housed in colonies (Figure 7D). While our findings do not demonstrate a difference in lifespan across groups, these results do show that we are more likely to find age-related changes reminiscent of those found in humans because they are studied at older ages than those from colonies.

### Domestic cats and wildcats share a common pace of development

We amassed observations (i.e., age of milestones) from wildcats (*Felis silvestris*) and domestic cats (*Felis catus*) to test whether domestication impacted the pace of development in cats (Figure 8A). We matched observations for statistics (e.g., minimum age of eye opening), sex, and environment (e.g., colony captive, captive, wild) and we excluded known cat breeds (e.g., Persian cats) for these comparisons. Observations are extracted from reproductive, behavioral and tooth development from both wild cats and non-breed domestic cats (Figure 8B). According to these data, the pace of development in domestic cats is similar to wildcats (Figure 8A). The slope of the linear model fit to the log-transformed age (in years post-conception) using domestic cats as the predictor and wild cats as the dependent variable (y=1.02+0.008; adj R^2^=0.97; p<0.01, df=154) is not significantly different from 1 (slope=1.02; SE=0.014; t=1.43; df=154, p=0.155; Figure 8A). Our findings therefore suggest that there was little change in the pace of development in the domestication of non-breed domesticated cats.

**Figure 8.**
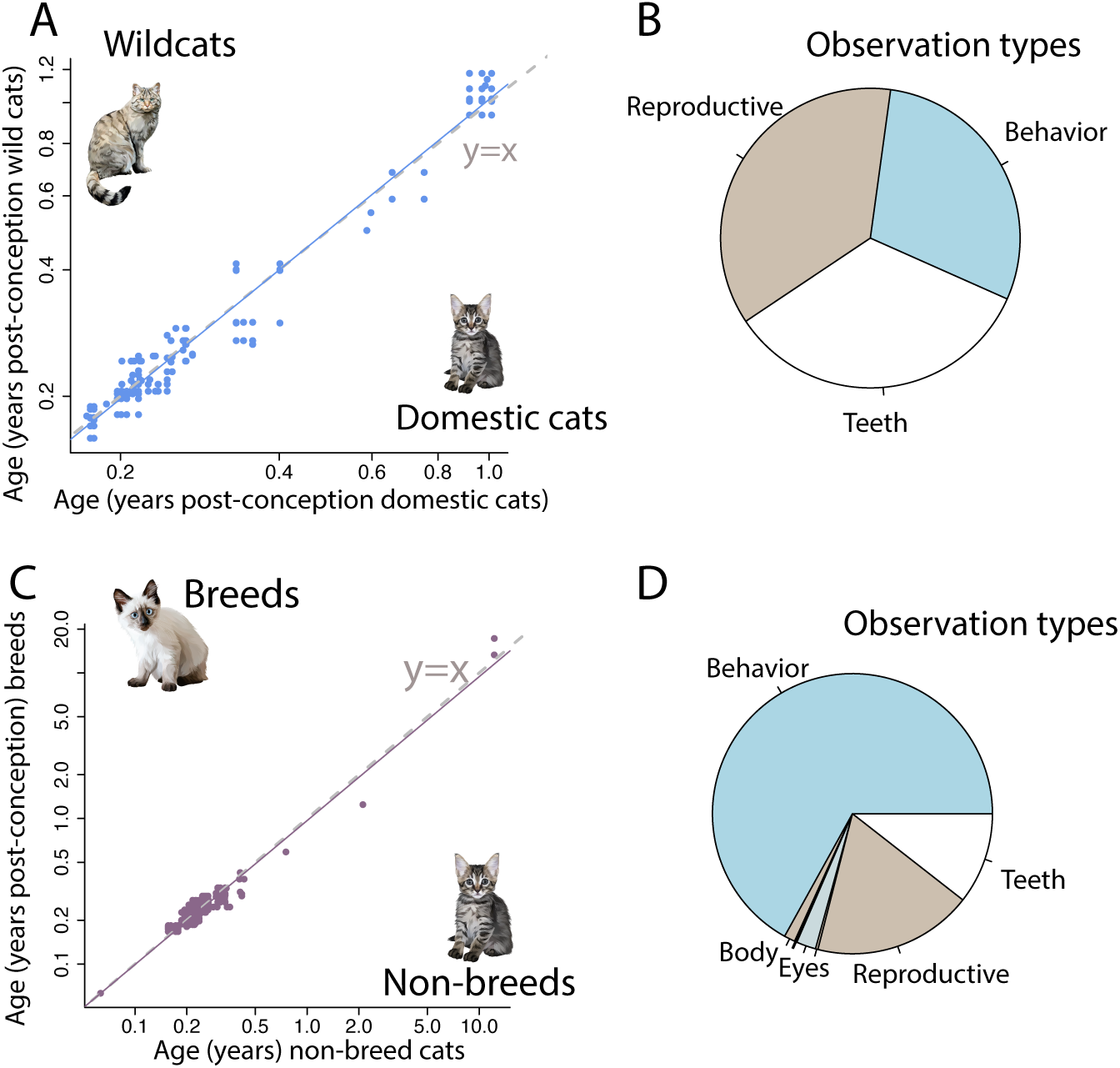
(A) Domestic cats and wildcats mature at similar rates. The fit to the model has a slope close to 1 (y=x). Observations (e.g., age of eye opening) were matched for sex, environment, and statistics (e.g., minimum age of eye opening, maximum of eye opening). (B) These observations capture the pace of behavioral, reproductive, and tooth development. (C) Breeding does not necessarily lead to modifications in the pace of development. A linear model fit to breed, and non-breed cats shows that the linear model is very close to 1. (D) These observations primarily capture behavioral, reproductive and tooth maturation.

### Breeding is not necessarily associated with modifications in the pace of development

Overall, the pace of development between breeds and non-breed cats (including wildcats) appears similar (Figure 8B). We matched observations (e.g., age of eye opening), environment (e.g., captive colony captive client, wild), sex, and statistics (e.g., minimum versus maximum age of eye opening). Breed cats show no significant differences in the pace of development compared with non-breed cats. The majority of observations are from tooth This is evident because the slope of the linear model fit to observations (in log-transformed years post-conception) for non-breed cats as the predictor and age of observation in breed cats as the dependent variable (y=0.99x-0.01, adj R^2^=0.93; df=709; Figure 8B) is significantly different than 1 (slope: 0.98; SE=0.01; t stat=-1.07; p=0.29).

### Translating ages shows cats live to age like a human octogenarian

We fit a general linear model with a smooth spline basis expansion to enable a flexible fit of the data to the event scale (Figure 1A). We equate ages across species, including time points from age-related changes in brain structure, blood chemistry profiles, and behavioral milestones (Figure 2A). In some cases, we found multiple observations per statistics, sex, and species. We averaged observations per species, sex, and statistics. We then generated an event scale (ordering of time points) averaged from imputed time points available across species. Each time point is subtracted from the age of the earliest time point, and each time point is divided by the difference between the maximum and minimum values (Figure 1A). Early time points are assigned a score close to 0 and later time points are assigned a score close to 1. We fit a general linear model with the event scale as the independent variable and observations as the dependent variable. The model allows for relative accelerations or decelerations in the pace of development and aging across species (Figure 1A).

The model accounts for a significant and high percentage of the variance (F=3,905, p<0.01, adj R^2^=0.977; df=2,077). and equates ages across the lifespan of humans, chimpanzees, mice, and cats. According to this model, the pace of development and aging follow complex trajectories across species. The pace of postnatal development being stretched (unusually long) in both humans and chimpanzees compared with cats and mice (Figure 1A). We also found that not every animal lives to the equivalent of a human octogenarian (Figure 1G). Specifically, we found that humans in their 80s map onto cats in their mid-teens, but this is not the case for mice and chimpanzees. Scatterplots illustrate age equivalencies between pairs of species (Figure 2). Chimpanzees in their 40s map onto humans in their 40s and 50s. Mice around 1.5 years of age map onto humans in their mid to late 40s. These findings highlight that not every animal lives to the equivalent of a human octogenarian and that cats are useful model systems for human aging.

## Discussion

We used observations from multiple scales (e.g., blood, brain structure, pathologies) to compare the pace of development and aging across and within species. We focused on health metrics because we can study many individuals to encapsulate within-species variation in cross-species age alignments. We used MRI scans from clinical and research settings to study cats varying widely in age. Pet cats visiting the clinic are older than those from colonies, and they show age-related changes in brain structure that mirror those patterns found in humans. We primarily studied domestic shorthair pets, which may be spared from health problems found in cat breeds. We also discuss how our findings can improve human health via a One Health framework, which advances the notion that improving the health of animals through study can improve human health.

### Cats and humans share age-related changes in brain structure

We used multiple MR scanners from clinical and research settings to compare the pace of development and aging from cats in the clinic and those from the colony. We found that humans and cats share similar age-related changes in volume and other brain features (e.g., gyrification, interthalamic adhesion thickness). The brain and concomitant structures atrophy with age in humans. Cats also develop cerebral cortex plaques and tangles in their teens (i.e., 14.1 years post-birth; de Sousa et al., 2023), which are hallmarks of Alzheimer’s disease (Lorena Sordo et al., 2021; de Sousa, et al., 2023). Our findings point to commonalities in brain changes during normal aging and diseases of aging. We did not focus on behavioral changes in aging in cats because behavioral changes capturing cognitive dysfunction are not easily comparable between humans and companion animals. There are currently no well-established methods to study cognitive dysfunctions in cats (Gunn-Moore et al., 2007; Gunn-Moore, 2011). We did not find evidence of cognitive dysfunction in cats receiving MRI scans. Cats with suspected dementia may not be scanned in clinical settings perhaps because there is no cure for cognitive dysfunction and risking the health of aged animals with anesthesia for the MRI is not warranted. Instead, cats typically have their brains scanned for acute reasons (e.g., epilepsy), rather than for health problems that build gradually (e.g., cognitive dysfunction). We therefore consider we studied humans and cats without any clear signs of cognitive dysfunction and that age-related changes in brain structure found in humans and cats reflect a healthy aging.

### Translating Time shows cats live to the equivalent of a human octogenarian

We relied on multiple metrics to translate ages across the lifespan of species (Figure 1). Our data shows that 80-year-old humans map onto cats in their teens, though not every animal lives to the equivalent of a human octogenarian. For example, 40-year-old chimpanzees map onto humans in their 50s (Charvet, 2021; Charvet et al., 2023). Very few chimpanzees have ages that map to human octogenarians because many do not live beyond their 40s. Accordingly, very few chimpanzees show signs of brain atrophy, plaques and tangles, and menopause because so few of them live to ages where such processes should emerge. A similar situation is observed in mice where a 2-year-old mouse maps onto a human in their 70s so that very few of them should develop brain atrophy, and other age-related changes. Our study shows that cats, and especially pet cats, live to be equivalent to a human octogenarian.

### Pet cats live to the equivalent of a human octogenarian

We found that pet cats’ somatic maturation was significantly longer to mature than colony cats, though such changes were not clearly observed from comparative MRI analyses. Species differences in the pace of development hold across four populations (two pet groups, two colony groups). Several possibilities exist as to why colony cats mature faster than pet cats. Colony cats may be bred and selected by humans to mature rapidly. Alternatively, inbreeding in colony cats may lead to faster rates of development compared with pets. Whereas the reproduction of colony cats is under human control, the reproduction of domestic shorthairs is not typically under human control. In many areas of the US, cats tend to reproduce in overabundance and with little human control. Genetic differences between colony and pet cats might impact the pace of development, though we can’t discount the possible importance of the environment in dictating developmental differences between pet and colony cats.

### Human and chimpanzees’ postnatal development is extended compared with cats and mice

We found that certain phases within the lifespan are relatively stretched in some species relative to others. Specifically, we found that humans possess a relatively extended postnatal period. Childhood and adolescence are phases that are relatively extended compared with cats, mice, and rats. Interestingly, postnatal development in humans is not noticeably extended compared with chimpanzees (Charvet, 2021; Charvet et al., 2023), suggesting that the relative extension in the duration of postnatal development evolved before the emergence of the modern human lineage. The pace of development and aging shows a complex relationship. Multiplying a species’ age by a factor is not a satisfactory means to equate ages across species.

### Aged pet cats inform human aging

One key finding from this present work is that pet cats are studied at significantly older ages than those housed in colonies. This holds true when comparing the ages of cat populations for which blood chemistry profiles and MRIs were collected. While these observations, by themselves, do not mean that the lifespan of pet cats is different from those housed in colonies, they do show that there is a natural propensity to study older cats in clinical settings rather than in a laboratory. Studying aged animals in a laboratory is challenging because cats need to be housed in colonies for a significant amount of time (e.g., 15 years) before they become aged. Pet cats represent a viable group of animals to study aging, especially considering that owners are sponsoring expensive diagnostics (e.g., MRIs) with the aim of diagnosing and treating diseases. Most of our work is focused on the domestic shorthair, which may outlive some cat and dog breeds (Sánchez-Vizcaíno et al., 2017; Mata, 2025), and pet domestic shorthairs as potential models to understand human aging.

### One Health: a framework to understand human health

We propose broadening how we study animals to understand health. We have overwhelmingly studied mice in a laboratory setting to study human diseases even though many of these mice do not spontaneously develop human diseases, and they are genetically engineered to artificially recreate aspects of human diseases. Findings from models often fail to translate to humans because artificial recreations of a human disease fail to generate meaningful translations for human diseases. We propose to increase focus on animals that spontaneously develop diseases akin to those found in humans. Cats offer a unique opportunity for translational research, as they share human environments, manifest similar disease processes (e.g., obesity, diabetes, arthritis), and undergo age-related patterns also found in humans. Experiments can aim to improve conditions and lifespan in animals. For example, there are positive and relatively non-invasive interventions (e.g., exercise) researchers could explore to improve cognition and lifespan in cats Studies that apply a One Health approach have a proven track record of enhancing the health of animals and humans (CDC, 2015). The study of a spontaneous genetic mutation causing GM1 gangliosidosis, a fatal enzyme disorder in both cats and humans, highlights the potential for veterinary research to advance human medicine. Some cats naturally develop this disease, and researchers have synthesized an interventional gene therapy that delivers a functional copy of the defective gene. Results in cats showed improved enzymatic activity and neuromuscular function while extending survival (McCoy, 2019). These promising preclinical findings led to the launch of human clinical trials, which produced encouraging results as children previously unable to walk or eat now can (Talesnik, 2022; Martin, 2022). Broadening our definition of model systems to incorporate a One Health approach and one that targets positive change in the health of cats can work to make significant strides in improving human health.

## Acknowledgments

We used data from the Primate Aging Database (https://primatedatabase.org/), which is an initiative of the National Institute on Aging.

---

**Table S1.**
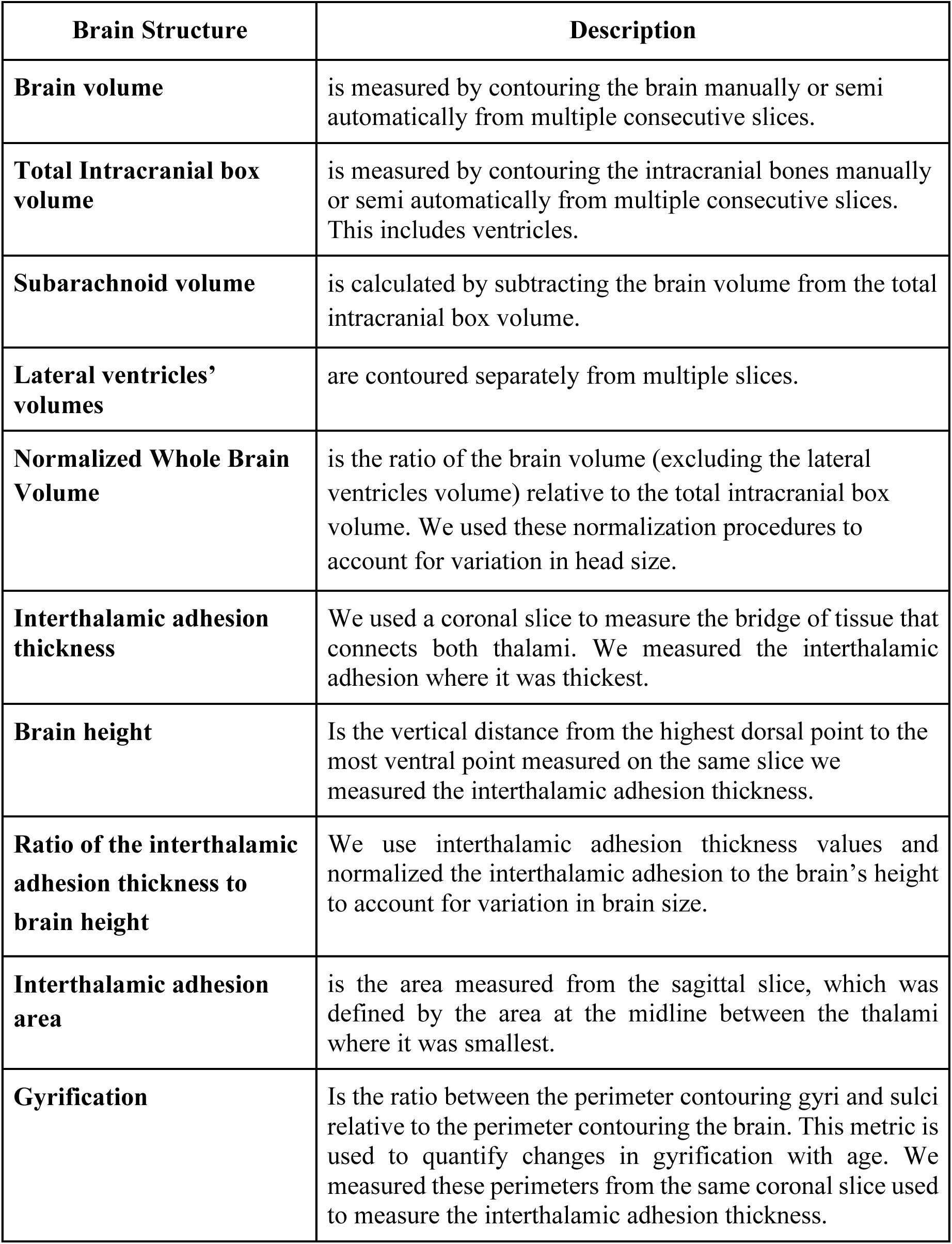
Description of metrics used to capture age-related changes in brain structure.

| **Brain Structure** | **Description** |
| --- | --- |
| **Brain volume** | is measured by contouring the brain manually or semi automatically from multiple consecutive slices. |
| **Total Intracranial box volume** | is measured by contouring the intracranial bones manually or semi automatically from multiple consecutive slices. This includes ventricles. |
| **Subarachnoid volume** | is calculated by subtracting the brain volume from the total intracranial box volume. |
| **Lateral ventricles' volumes** | are contoured separately from multiple slices. |
| **Normalized Whole Brain Volume** | is the ratio of the brain volume (excluding the lateral ventricles volume) relative to the total intracranial box volume. We used these normalization procedures to account for variation in head size. |
| **Interthalamic adhesion thickness** | We used a coronal slice to measure the bridge of tissue that connects both thalami. We measured the interthalamic adhesion where it was thickest. |
| **Brain height** | Is the vertical distance from the highest dorsal point to the most ventral point measured on the same slice we measured the interthalamic adhesion thickness. |
| **Ratio of the interthalamic adhesion thickness to brain height** | We use interthalamic adhesion thickness values and normalized the interthalamic adhesion to the brain's height to account for variation in brain size. |
| **Interthalamic adhesion area** | is the area measured from the sagittal slice, which was defined by the area at the midline between the thalami where it was smallest. |
| **Gyrification** | Is the ratio between the perimeter contouring gyri and sulci relative to the perimeter contouring the brain. This metric is used to quantify changes in gyrification with age. We measured these perimeters from the same coronal slice used to measure the interthalamic adhesion thickness. |

## References

Berg L, McKeel DW, Miller JP, Storandt M, Rubin EH, Morris JC, Baty J, Coats M, Norton J, Goate AM, Price JL (1998) Clinicopathologic studies in cognitively healthy aging and Alzheimer disease: relation of histologic markers to dementia severity, age, sex, and apolipoprotein E genotype. Archives of neurology. 55:326–35.

Chambers JK, Tokuda T, Uchida K, Ishii R, Tatebe H, Takahashi E, Tomiyama T, Une Y, Nakayama H (2015) The domestic cat as a natural animal model of Alzheimer’s disease. Acta Neuropathol Commun 3:78.

Charvet CJ (2021) Cutting across structural and transcriptomic scales translates time across the lifespan in humans and chimpanzees. Proc Biol Sci 288:20202987.

Charvet CJ, Ofori K, Baucum C, Sun J, Modrell MS, Hekmatyar K, Edlow BL, van der Kouwe AJ (2022) Tracing Modification to Cortical Circuits in Human and Nonhuman Primates from High-Resolution Tractography, Transcription, and Temporal Dimensions. J Neurosci. 42:3749–3767.

Charvet CJ, de Sousa AA, Vassilopoulos T (2025) Translating time: challenges, progress, and future directions. Brain Res Bull 221:111212.

Charvet CJ, Finlay BL (2018) Comparing adult hippocampal neurogenesis across species: translating time to predict the tempo in humans. Front Neurosci 12:706.

Charvet CJ, Ofori K, Falcone C, Rigby Dames BA (2023) Transcription, structure, and organoids translate time across the lifespan of humans and great apes. PNAS Nexus 2:pgad230.

Chen X, Errangi B, Li L, Glasser MF, Westlye LT, Fjell AM, Walhovd KB, Hu X, Herndon JG, Preuss TM, Rilling JK (2013) Brain aging in humans, chimpanzees (*Pan troglodytes*), and rhesus macaques (*Macaca mulatta*): magnetic resonance imaging studies of macro- and microstructural changes. Neurobiol Aging 34:2248–2260.

Clancy B, Darlington RB, Finlay BL (2001) Translating developmental time across mammalian species. Neuroscience 105:7–17.

Cleary P, Shorvon S, Tallis R (2004) Late-onset seizures as a predictor of subsequent stroke. Lancet 363:1184–1186.

Cottam NC, Ofori K, Stoll KT, Bryant M, Rogge JR, Hekmatyar K, Sun J, Charvet CJ (2025) From circuits to lifespan: translating mouse and human timelines with neuroimaging-based tractography. J Neurosci 45:e1429242025.

Coupé P, Manjón JV, Lanuza E, Catheline G (2019) Lifespan changes of the human brain in Alzheimer’s disease. Sci Rep 9:3998.

Davidson YS, Robinson A, Prasher VP, Mann DMA (2018) The age of onset and evolution of Braak tangle stage and Thal amyloid pathology of Alzheimer’s disease in individuals with Down syndrome. Acta Neuropathol Commun 6:56.

de Sousa FG, Queiroz FFS, Muzzi RAL, Veado JCC, Beier SL (2025) Systemic arterial hypertension and factors associated with blood pressure dysregulation in companion animals. Vet Sci 12:453.

Driscoll CA, Clutton-Brock J, Kitchener AC, O’Brien SJ (2009a) The Taming of the Cat. Genetic and archaeological findings hint that wildcats became housecats earlier--and in a different place--than previously thought. Sci Am 300:68–75.

Driscoll CA, Macdonald DW, O’Brien SJ (2009b) From wild animals to domestic pets, an evolutionary view of domestication. Proc Natl Acad Sci USA 106:9971–9978.

Edler MK, Sherwood CC, Meindl RS, Hopkins WD, Ely JJ, Erwin JM, Mufson EJ, Hof PR, Raghanti MA (2017) Aged chimpanzees exhibit pathologic hallmarks of Alzheimer’s disease. Neurobiol Aging 59:107–120.

Finch CE, Austad SN (2015) Commentary: is Alzheimer’s disease uniquely human? Neurobiol Aging 36:553–555.

Fjell AM, McEvoy L, Holland D, Dale AM, Walhovd KB, Alzheimer’s Disease Neuroimaging Initiative (2013) Brain changes in older adults at very low risk for Alzheimer’s disease. J Neurosci 33:8237–8242.

Fotenos AF, Snyder AZ, Girton LE, Morris JC, Buckner RL (2005) Normative estimates of cross-sectional and longitudinal brain volume decline in aging and AD. Neurology 64:1032–1039.

Freeman LM, Abood SK, Fascetti AJ, Fleeman LM, Michel KE, Laflamme DP, Bauer C, Kemp BLE, Van Doren JR, Willoughby KN (2006) Disease prevalence among dogs and cats in the United States and Australia and proportions of dogs and cats that receive therapeutic diets or dietary supplements. J Am Vet Med Assoc 229:531–534.

Ginel PJ, Lucena R, Fernández M (2002) Duration of increased serum alkaline phosphatase activity in dogs receiving different glucocorticoid doses. Res Vet Sci 72:201–204.

Gunn-Moore DA (2011) Cognitive dysfunction in cats: clinical assessment and management. Top Companion Anim Med. 26(1):17–24.

Gray-Edwards HL, Salibi N, Josephson EM, Hudson JA, Cox NR, Randle AN, McCurdy VJ, Bradbury AM, Wilson DU, Beyers RJ, Denney TS, Martin DR (2014) High resolution MRI anatomy of the cat brain at 3 Tesla. J Neurosci Methods 227:10–17.

Griffin B, Baker HJ (2002) Domestic cats as laboratory animals. Lab Anim Med 2:459–482. doi: 10.1016/b978-012263951-7/50015-6.

Gunn-Moore D, Moffat K, Christie LA, Head E (2007) Cognitive dysfunction and the neurobiology of ageing in cats. J Small Anim Pract 48:546–553.

Hadjicharalambous C, Alpantaki K, Chatzinikolaidou M (2021) Effects of NSAIDs on pre-osteoblast viability and osteogenic differentiation. Exp Ther Med 22:740.

Hasegawa D, Yayoshi N, Fujita Y, Fujita M, Orima H (2005) Measurement of interthalamic adhesion thickness as a criteria for brain atrophy in dogs with and without cognitive dysfunction (dementia). Vet Radiol Ultrasound 46:452–457.

Hazenfratz M, Taylor SM (2018) Recurrent seizures in cats: diagnostic approach - when is it idiopathic epilepsy? J Feline Med Surg 20:811–823.

Knight A (2008) The beginning of the end for chimpanzee experiments? Philos Ethics Humanit Med.3:16. doi: 10.1186/1747-5341-3-16.

Kraska A, Dorieux O, Picq J-L, Petit F, Bourrin E, Chenu E, Volk A, Perret M, Hantraye P, Mestre-Frances N, Aujard F, Dhenain M (2011) Age-associated cerebral atrophy in mouse lemur primates. Neurobiol Aging 32:894–906.

Landsberg GM, Nichol J, Araujo JA (2012) Cognitive dysfunction syndrome: a disease of canine and feline brain aging. Vet Clin N Am Small Anim Pract 42:749–768.

Levy JK, Crawford PC, Werner LL (2006) Effect of age on reference intervals of serum biochemical values in kittens. J Am Vet Med Assoc 228:1033–1037.

Long X, Liao W, Jiang C, Liang D, Qiu B, Zhang L (2012) Healthy aging: an automatic analysis of global and regional morphological alterations of human brain. Acad Radiol 19:785–793.

Marcus DS, Wang TH, Parker J, Csernansky JG, Morris JC, Buckner RL (2007) Open access series of imaging studies (OASIS): cross-sectional MRI data in young, middle aged, nondemented, and demented older adults. J Cogn Neurosci 19:1498–1507.

Mata F (2025) Life expectancy of cats in Britain: moggies and mollies live longer. PeerJ 13:e18869.

Nakai N, Nawa K, Maekawa M, Nagasawa H (1992) Age-related changes in hematological and serum biochemical values in cats. Experimental Animals. 41:287–94.

Nelson PT, Braak H, Markesbery WR (2009) Neuropathology and cognitive impairment in Alzheimer disease: a complex but coherent relationship. J Neuropathol Exp Neurol 68:1–14.

Noh D, Choi S, Choi H, Lee Y, Lee K (2017) Evaluation of interthalamic adhesion size as an indicator of brain atrophy in dogs with and without cognitive dysfunction. Vet Radiol Ultrasound 58:581–587.

Oikonomidis IL, Milne E (2023) Clinical enzymology of the dog and cat. Aust Vet J 101:465–478.

Picq JL, Aujard F, Volk A, Dhenain M (2012) Age-related cerebral atrophy in nonhuman primates predicts cognitive impairments. Neurobiol Aging 33:1096–1109.

Pineda C, Aguilera-Tejero E, Guerrero F, Raya AI, Rodriguez M, Lopez I (2013) Mineral metabolism in growing cats: changes in the values of blood parameters with age. Journal of feline medicine and surgery.15(10):866–71.

Richards J (2019) Neurodevelopmental MRI database [Tool/Resource.]. Washington: NITRC.

Rofina JE, van Ederen AM, Toussaint MJM, Secrève M, van der Spek A, van der Meer I, Van Eerdenburg FJCM, Gruys E (2006) Cognitive disturbances in old dogs suffering from the canine counterpart of Alzheimer’s disease. Brain Res 1069:216–226.

Rigby Dames BAR, Kilili H, Charvet CJ, Díaz-Barba K, Proulx MJ, de Sousa AA, Urrutia AO (2023) Evolutionary and genomic perspectives of brain aging and neurodegenerative diseases. Prog Brain Res 275:165–215.

Rosen RF, Farberg AS, Gearing M, Dooyema J, Long PM, Anderson DC, Davis-Turak J, Coppola G, Geschwind DH, Paré J-F, Duong TQ, Hopkins WD, Preuss TM, Walker LC (2008) Tauopathy with paired helical filaments in an aged chimpanzee. J Comp Neurol 509:259–270.

Syme HM, Barber PJ, Markwell PJ, Elliott J (2002) Prevalence of systolic hypertension in cats with chronic renal failure at initial evaluation. J Am Vet Med Assoc. 220:1799–804.

Sordo L, Martini AC, Houston EF, Head E, Gunn-Moore D (2021) Neuropathology of aging in cats and its similarities to human Alzheimer’s disease. Front Aging 2:684607.

Sánchez-Vizcaíno F, Noble PJM, Jones PH, Menacere T, Buchan I, Reynolds S, Dawson S, Gaskell RM, Everitt S, Radford AD (2017) Demographics of dogs, cats, and rabbits attending veterinary practices in Great Britain as recorded in their electronic health records. BMC Vet Res 13:218.

Seibert L (2017) Management of dogs and cats with cognitive dysfunction. Today’s Vet Pr 7:1– 8.

Sordo L, Breheny C, Halls V, Cotter A, Tørnqvist-Johnsen C, Caney SMA, Gunn-Moore DA (2020) Prevalence of disease and age-related behavioural changes in cats: past and present. Vet Sci 7:85.

Tacutu R, Craig T, Budovsky A, Wuttke D, Lehmann G, Taranukha D, Costa J, Fraifeld VE, de Magalhães JP (2013) Human ageing genomic resources: integrated databases and tools for the biology and genetics of ageing. Nucleic Acids Res 41:D1027–D1033.

Teng KTY, Brodbelt DC, Church DB, O’Neill DG (2024) Life tables of annual life expectancy and risk factors for mortality in cats in the UK. J Feline Med Surg 26:1098612X241234556.

Vaden SL, Pressler BM, Lappin MR, Jensen WA (2004) Effects of urinary tract inflammation and sample blood contamination on urine albumin and total protein concentrations in canine urine samples. Vet Clin Pathol 33:14–19.

Vaske HH, Schermerhorn T, Grauer GF (2016) Effects of feline hyperthyroidism on kidney function: a review. J Feline Med Surg 18:55–59.

Vervloet MG, Sezer S, Massy ZA, Johansson L, Cozzolino M, Fouque D (2017) The role of phosphate in kidney disease. Nat Rev Nephrol 13:27–38.

Workman AD, Charvet CJ, Clancy B, Darlington RB, Finlay BL (2013) Modeling transformations of neurodevelopmental sequences across mammalian species. J Neurosci. 33:7368–83.

Yokoyama M, Kobayashi H, Tatsumi L, Tomita T (2022) Mouse models of Alzheimer’s disease. Front Mol Neurosci 15:912995.

Youssef SA, Capucchio MT, Rofina JE, Chambers JK, Uchida K, Nakayama H, Head E (2016) Pathology of the aging brain in domestic and laboratory animals, and animal models of human neurodegenerative diseases. Vet Pathol 53:327–348.

